# Percent amplitude of fluctuation: a simple measure for resting-state fMRI signal at single voxel level

**DOI:** 10.1101/214098

**Authors:** Xi-Ze Jia, Gong-Jun Ji, Wei Liao, Ya-Ting Lv, Jue Wang, Ze Wang, Han Zhang, Dong-Qiang Liu, Yu-Feng Zang

**Author notes:** Corresponding to ZANG Yu-Feng, M.D. Center for Cognition and Brain Disorders Hangzhou Normal University, Room 261, Building 7, Affiliated Hospital of Hangzhou Normal University No. 126, Wenzhou Rd, Hangzhou, Zhejiang, 310015, China.

## Abstract

The amplitude of low-frequency fluctuation (ALFF) measures resting-state functional magnetic resonance imaging (RS-fMRI) signal of each voxel. However, the unit of blood oxygenation level-dependent (BOLD) signal is arbitrary and hence ALFF is sensitive to the scale of raw signal. A well-accepted standardization procedure is to divide each voxel’s ALFF by the global mean ALFF. However, this makes the individual voxel’s ALFF dependent on the global mean. Although Fractional ALFF (fALFF), proposed as a ratio of the ALFF to the total amplitude within the full frequency band, offers possible solution of the standardization, it actually mixes with the fluctuation power within the full frequency band and thus cannot reveal the true amplitude characteristics of a given frequency band. We proposed a new standardized, stand-alone, single-voxel metrics for RS-fMRI, namely percent amplitude of fluctuation (PerAF). PerAF is an analog to the percent signal change that has been widely used in the task fMRI communities, which allows it to be a straightforward measurement of BOLD signal fluctuations during resting state. We further conducted a test-retest reliability analysis comparing the relevant metrics, which indicated that PerAF was generally more reliable than the ALFF and fALFF. In a real RS-fMRI application, we further demonstrated that with and without standardization by global mean PerAF yielded prominently different results when comparing eyes open with eyes closed resting conditions, suggesting that future study should provide both with and without global mean standardization. The above results suggest that PerAF is a more reliable, straightforward and promising measurement for voxelwise brain activity-based RS-fMRI studies. For prompting future application of PerAF, we also implemented this method into a user-friendly toolbox *REST-PerAF*.

## 1. Introduction

The neuroimaging field has an increasing research interest in assessing the spontaneous brain activity using BOLD RS-fMRI. Functional connectivity (FC), first used in a RS-fMRI study more than two decade ago (Biswal et al. 1995), is still among the most widely used methods in the RS-fMRI communities. However, by focusing on temporal synchronization of the BOLD signals between any pair of brain regions, FC analysis cannot infer any information of the regional spontaneous brain activity. Another widely used RS-fMRI measurement, ALFF (Zang et al. 2007), can be adopted on this purpose, as it provides direct characterization to spontaneous brain activity at each voxel. However, since BOLD signal has arbitrary units, ALFF is sensitive to the scale of the raw BOLD signal and cannot be directly fed into following statistical analysis. An approach to deal with such a scaling effect is to normalize the raw ALFF value by the global mean ALFF, the averaged ALFF value across all voxels in the brain (Yan et al. 2013b; Zang et al. 2007; Zuo et al. 2010a). However, such manipulation will make the voxelwise ALFF depend on the global mean. In an extreme case, certain brain lesions may significantly alter the mean intensity of the BOLD signal. This can have significant effect on the derived global mean of the ALFF and therefore lead a bias of the normalized ALFF. In some other ALFF studies with partial brain coverage where the global mean is not available (Lv et al. 2013; Yang et al. 2007; Yang et al. 2011; Yuan et al. 2014), to handle the scaling problem, an slightly modified version of global mean ALFF, the “mask mean” ALFF, has to be used to calibrate raw ALFF. In such cases, it is difficult to compare the results from different studies, because it is hard to use the same mask for calcluating the “mask mean” ALFF for calibrating raw ALFF.

As an another voxelwise measurement and a derivative from ALFF, fractional ALFF (fALFF) was proposed to normalize ALFF. The fALFF is a ratio of the ALFF within a specific low frequency band (usually 0.01 - 0.08 Hz) to the total BOLD fluctuation amplitude within the full frequency band (Zou et al. 2008). It can be regarded as a standardized ALFF-like metric at the single voxel level and is theoretically a scale-independent (i.e., not depending on the mean absolute value of the BOLD signals) method. One-sample t-test on fALFF within a group of participants did confirm that a much better default mode network pattern could be captured by fALFF (i.e., significantly higher fALFF within the default mode network) compared to ALFF (Zou et al. 2008). Previous study have also shown that the fALFF measured in resting state can significantly reduce inter-subject variability of task fMRI activations (Kalcher et al. 2013). However, fALFF is a mixture of frequency-specific fluctuation amplitude and whole-frequency-band fluctuation amplitude. In other words, it depends on both values, and cannot reveal the true amplitude characteristics of a given frequency band per se. Given different voxels may have different whole-frequency-band fluctuation amplitude, the resultant fALFF could be highly dominated by the changing whole-frequency-band fluctuation amplitude from voxel to voxel. For example, if one is interested in characterizing the fluctuation amplitude of the whole frequency band by using fALFF, the result will always be one (i.e., the ALFF value divided by itself according to the definition of fALFF). Moreover, previous studies have shown that fALFF had generally lower test-retest reliability than that of ALFF in gray matter voxels (Zuo et al. 2010a; Küblböck et al. 2014). All the evidences questioning whether fALFF is a good metric for RS-fMRI. We think that this problem is getting more and more serious because of the researches trend focusing on specific frequency band. Recently, frequency-dependent or frequency-specific analysis of RS-fMRI has been drawing increasing attention (Han et al. 2011; Wee et al. 2012; Esposito et al. 2013; Wei et al. 2014; Huang et al. 2014; Yu et al. 2014; Yue et al. 2014); many studies have suggested that frequency-specific BOLD fluctuations can be used to detect disease biomarkers (Malinen et al. 2010; Otti et al. 2013) and to detect different subject’s status (Yuan et al. 2014). From the above analysis, fALFF may not be a proper standardized metric for these studies.

In task fMRI studies, percent signal change is a popular measurement of task-induced BOLD signal changes, which measures the difference in fMRI signal between the baseline condition (B) and the task condition (T), i.e., percent signal change = (T-B)/B×100%. Typical percent signal change is approximately 1% - 3% in block design (Kanwisher et al. 1997; Kanwisher et al. 1998) and from 0.2% to 1% in event-related design (Grill-Spector et al. 2004; Tambini et al. 2010). Although RS-fMRI data has no such explicit task vs. control design, a similar metric to the percent signal change can be formulated for RS-fMRI by measuring the percentage of BOLD fluctuations relative to the mean BOLD signal intensity for each time point and averaging across the whole time series, namely “Percentage Amplitude Fluctuation”, short for *PerAF*. As compared with ALFF, PerAF is a scale-independent method. PerAF can also aviod from the confounding mixture from voxel-specific fluctuation amplitude in fALFF. Hence, PerAF seems to be a promising metric of voxelwise spontaneous BOLD activity.

Next, we firstly provide detailed formation of PerAF, and then test its validity with simulated data. We further assess the test-retest reliability of PerAF using a real RS-fMRI data. At last, we demonstrated the feasibility of using PerAF to detect voxelwise differences between two resting states (eyes open [EO] vs. eyes closed [EC]) in another dataset. To facilitate future applications, we implemented PerAF as a graph user interface (GUI)-based function into an existing widely-adopted MATLAB-based Resting-state fMRI data analysis Toolkit (REST) (Song et al. 2011) and Statistical Parametric Mapping (SPM) (Friston et al. 1994). We also made a command-line-based function of PerAF for calculating this metric in LINUX based on Analysis of Functional Neuro-Images (AFNI) (Cox 1996).

## 2. Methods and Results

### 2.1. PerAF calculation and experiment on simulated data

The PerAF of each voxel was calculated as follows,

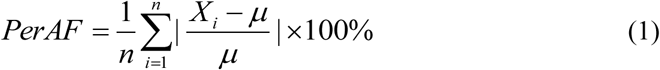

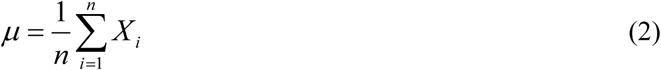

where *X*_*i*_ is the signal intensity of the *i*_*th*_ time point, *n* is the total number of time points of the time series, and *μ* is the mean value of the time series.

A simulated time series *X*_*1*_ was created, which contained 120 time points with random values. Its derivative time series *X*_*2*_ and *X*_*3*_ were generated by multiplying *X*_*1*_ by *2* and by 3, respectively (Fig. 1). PerAF were calculated for each time series. For comparison purpose, we also calculated ALFF (see Section “2.2.3” for detailed ALFF calculation). As shown in Fig. 1, the ALFF value is proportional to the mean value of the time series, but PerAF is not. Because the absolute BOLD signal intensity has arbitrary units, ALFF results will be effected by the scale of BOLD signal. The simulated data showed that, without a standardization procedure, ALFF can not be used for direct comparison.

**Fig. 1.**
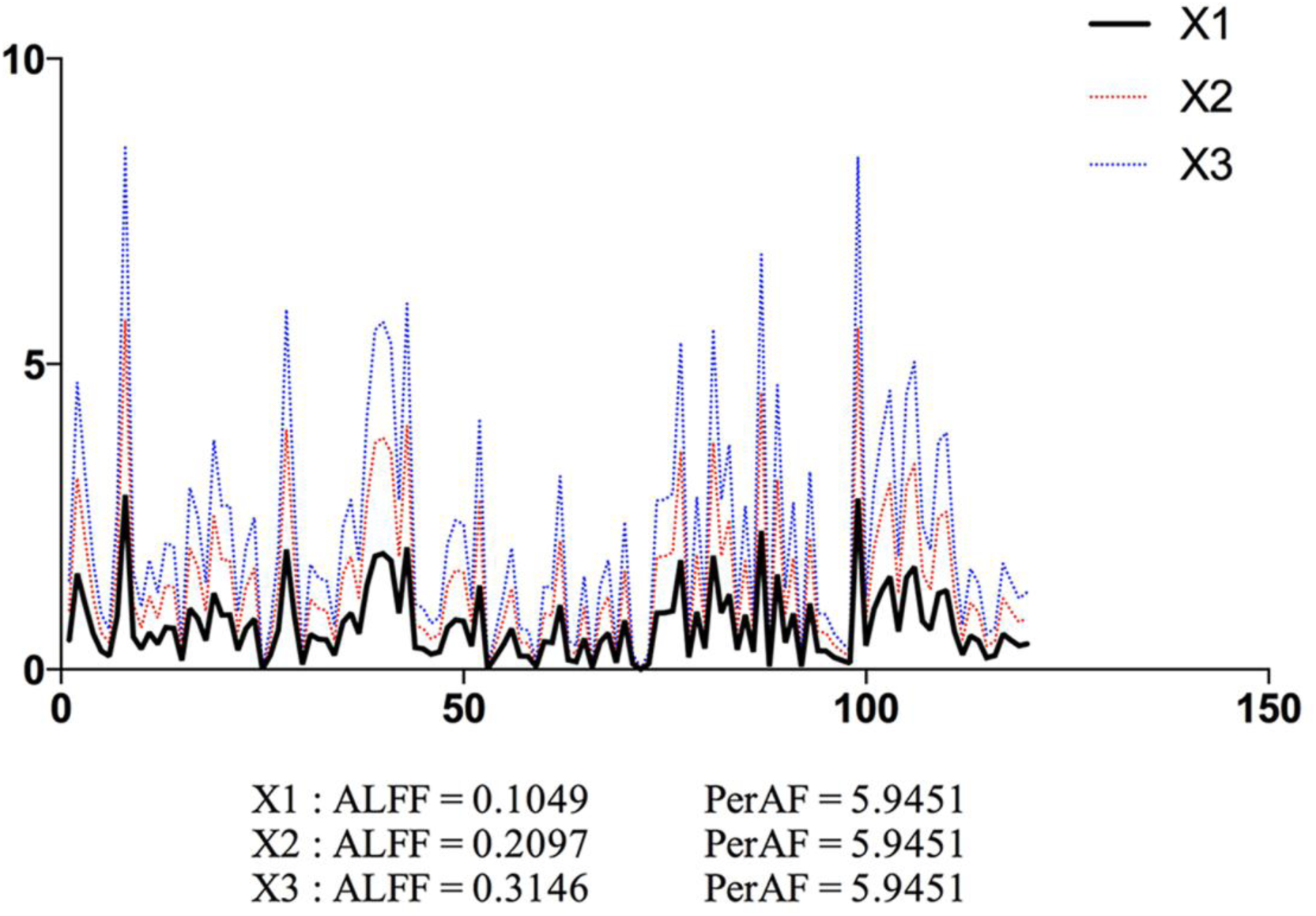
Simulated “resting-state” BOLD time series. The ALFF and PerAF of a simulated time series (X1), and its derivative time series of multiplying A by 2 (X2) and by 3 (X3). ALFF and PerAF were calculated for each time series. ALFF value is proportional to the mean value of the time series, but PerAF keeps the same. X and y axis are time and intensity, respectively, in arbitrary unit.

### 2.2. Dataset-1: test-retest reliability assessment

#### 2.2.1. Data description

Dataset-1 was from the International Neuroimaging Data-Sharing Intiative (INDI) (http://www.nitrc.org/projects/nyu_trt). This dataset includes 25 participants (mean age 30.7 ± 8.8 years, 16 females). Participants did not have any history of psychiatric or neurological illness. Informed consent was obtained from all participants. The study was approved by the Institutional Review Board of the New York University School of Medicine and New York University. Participants had three resting state sessions. Session 2 and 3 were collected 45 min apart, and were 5 - 16 months (mean 11 ± 4 months) after Session 1. During each scanning session, participants were instructed to continuously keep eyes open and a word “Relax” was centrally projected in white against a black background.

Dataset-1 was obtained using a 3T Siemens (Allegra) scanner. Each scan consisted of 197 contiguous EPI functional volumes (TR = 2000 ms; TE = 25 ms; flip angle = 90°; 39 axial slices; field of view (FOV) = 192 × 192 mm^2^; matrix = 64 × 64; acquisition voxel size = 3 × 3 × 3 mm^3^). For more information regarding Dataset-1 collection, please refer to (Shehzad et al. 2009).

#### 2.2.2. Data preprocessing

The preprocessing was performed using Data Processing Assistant for Resting-State fMRI (DPARSF) (Chao-Gan and Yu-Feng 2010) (http://www.restfmri.net), including: 1) discarding the first 10 timepoints for the longitudinal magnetization to reach a steady state and for participant’s adaptation to the scanning noise; 2) slice timing correction; 3) head motion correction; 4) co-registration, spatial normalization and resampling to 3 mm isotropic voxel size; 5) spatial smoothing with an isotropic Gaussian kernel with a FWHM of 6 mm; 6) removing the linear trend of the time series; and 7) band-pass (0.01-0.08Hz) filtering. Three participants were excluded from further analyses because of excessive head motion (more than 2.0 mm of maximal translation or 2.0^°^ of maximal rotation) throughout the course of scanning.

Considering the fact that head motion regression is drawing more and more attention in the RS-fMRI studies, we also explored the effect of head motion regression. Before band-pass filtering, we regressed Friston-24 head motion parameters individually. Friston-24 head motion parameters includes 6 head motion parameters (3 for transition and 3 for rotation), their historical effects (position in the previous scan, 6 parameters), and square of the 12 parameters (Friston et al. 1996). A recent RS-fMRI study comprehensively investigated the effects of a set of head motion parameters on a set of measurements and concluded that Friston-24 performed the best on most RS-fMRI measurements (Yan et al. 2013a).

#### 2.2.3. Test-retest reliability of PerAF, ALFF, and fALFF

The PerAF was calculated in the way as mentioned in section “2.1”. The ALFF and fALFF analysis was performed using REST (Song et al. 2011) (http://www.restfmri.net). After preprocessing, the time series for each voxel was transformed into the frequency domain with a fast Fourier transform (FFT) and the power spectrum was then obtained. Since the power of a given frequency is proportional to the square of the amplitude of this frequency component, the square root was calculated at each frequency of the power spectrum and the averaged square root was obtained across 0.01 - 0.08 Hz at each voxel. This averaged square root was taken as the ALFF (Zang et al. 2007). Then a ratio of the sum of amplitude within the low frequency band (i.e., ALFF) to that of the whole frequency band was computed as fALFF (Zou et al. 2008).

The original ALFF value is not very suitable for comparison, so ALFF of each voxel was divided by the global mean ALFF of each participant (Zang et al. 2007) (we call this result mALFF). The same procedure was performed for fALFF (Zou et al. 2008) (mfALFF). Mathematically, it is not necessary in PerAF since the scaling factor has been normalized by dividing the temporal mean. However, to have a fair comparison with ALFF and fALFF and to investigate the effect of global mean value, the PerAF of each voxel was also divided by the global mean PerAF of each participant (thus we have both PerAF and mPerAF). In the original paper reporting Dataset-1, the authors did not use the cerebellum and inferior part of temporal lobe because these brain areas in some participants were not covered (Shehzad et al. 2009). Therefore, we made an intersection mask within which all 75 scanning sessions (3 sessions for each of the 25 participants) were covered (Fig. 2). Specifically, the mean fMRI image of each session was spatially normalized and then binarized (using logical function from MATLAB). Then all 75 binary images and a whole brain mask which provided in the software REST (Song et al. 2011) were combine to the intersection mask. The following statistical analysis was constrained within this intersection mask. It has been proposed that z-transformation of ALFF could improve the normality of distribution acorss subjects (Zuo et al. 2010a). Therefore we also transformed the PerAF, ALFF, and fALFF to their respective z score maps, i.e., minus by global mean and divided by standard deviation (SD), thus generating zPerAF, zALFF, and zfALFF. The different metrics and their derivatives were summarized in Table 1. As the original ALFF value is not suitable for comparison, it was excluded from further analysis.

**Fig. 2.**
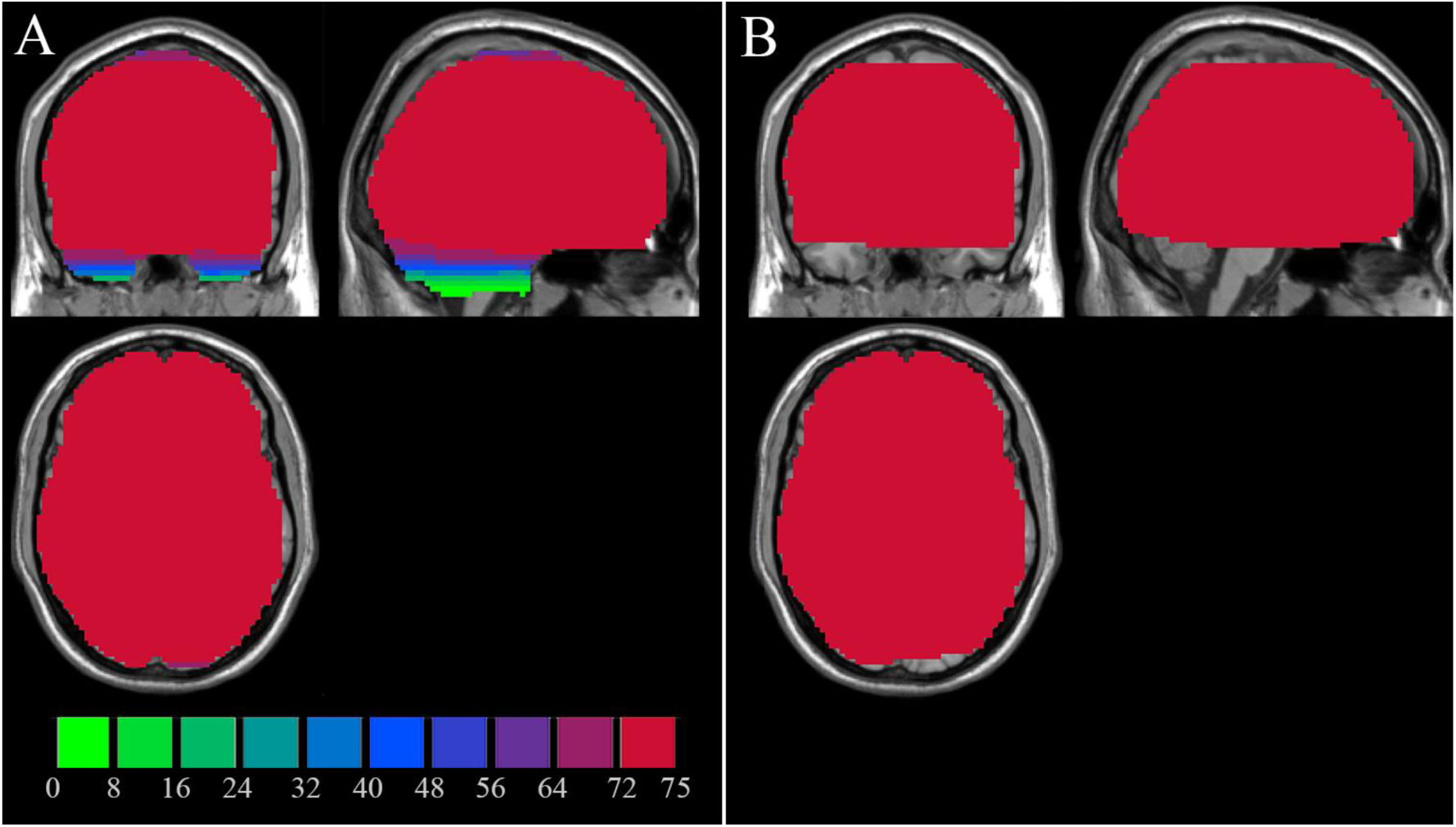
Intersection mask of Dataset-1. The left pannel shows how many sessions (totally 25 subjects × 3 = 75 sessions) were covered, for each voxel in Dataset-1. The right pannel is an intersection mask which was covered by all 75 sessions.

**Table 1.**
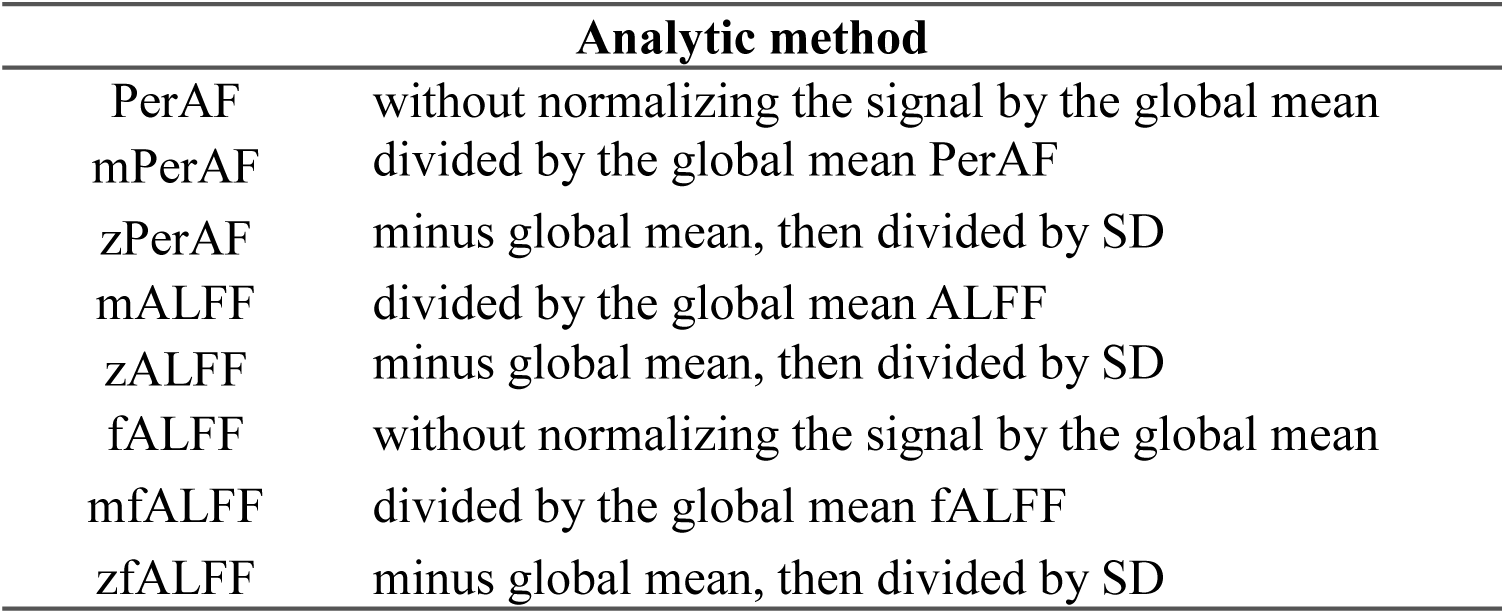
Standardization methods in the current study

To investigate the test-retest reliability of PerAF, ALFF, and fALFF over time, intraclass correlation coefficient (ICC) was calculated between each pair of the 3 sessions in Dataset-1. ICC has been widely used in previous studies for test-retest reliablility (Zuo et al. 2010a; Shehzad et al. 2009; Zuo et al. 2010b; Liao et al. 2013). Dataset-1 allows both long-term reliability (5 - 16 months apart) and short-term reliability (< 1h apart). ICC > 0.5 was considered as moderate or higher test-retest reliablility (Shehzad et al. 2009; Zuo et al. 2010a) and was used as a threshold in this study. As shown in Fig. 3, for all the measures including PerAF, mPerAF, zPerAF, mALFF, zALFF, fALFF, mfALFF, and zfALFF, most cortical areas showed moderate to high short-term (session 2 against session 3) test-retest reliability. Long-term test-retest reliability was lower than short-term reliability (See Fig. 3 for session 1 against session 2 and Supplementary Figure 1 (Fig. S1) for session 1 against session 3). Gray matter’s reliability was much higher compared to the white matter. fALFF and its derivative maps showed the worst test-retest reliability among the three metrics (Fig. 3).

**Fig. 3.**
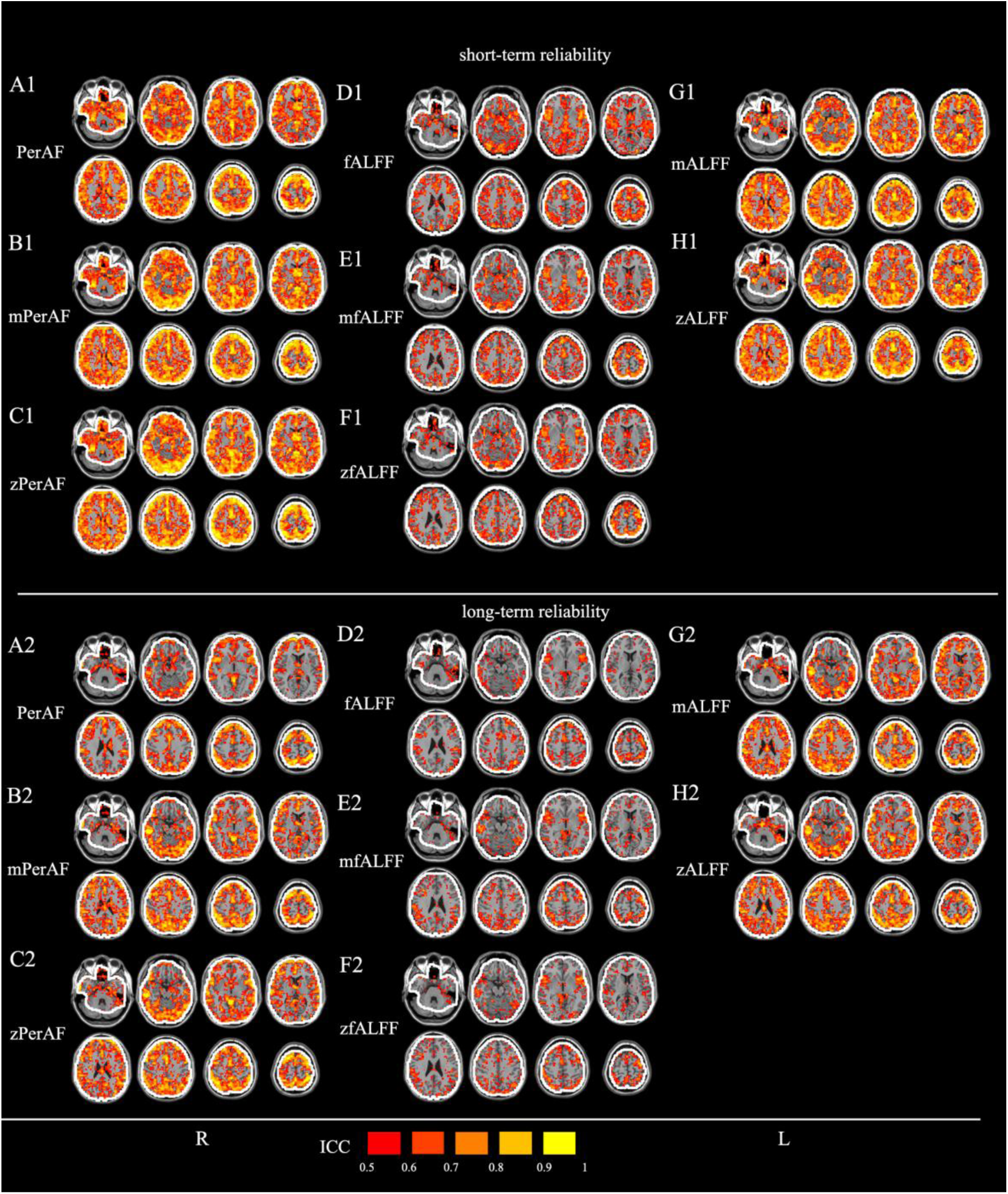
Test-retest reliability in Dataset-1. The upper part is for short-term (session 2 against session 3) (A1 - H1) and lower part is for long-term (session 1 against session 2) (A2 – H2). The intraclass correlation (ICC) maps include: (A1, A2) PerAF (without standardization by global mean), (B1, B2) mPerAF (divided by the global mean PerAF), (C1, C2) zPerAF (minus mean and divided by standard deviation), (D1, D2) fALFF (without standardization by global mean), (E1, E2) mfALFF (divided by the global mean fALFF), (F1, F2) zfALFF (minus mean and divided by standard deviation), (G1, G2) mALFF (divided by the global mean ALFF), and (H1, H2) zALFF (minus mean and divided by standard deviation). The original ALFF map is mathematically unsuitable for comparison and hence not listed here. The white contours denote the boundary of the intersection mask. The ICC threshold was set at ≥ 0.5 for all maps. L: left side of the brain. R: right side of the brain.

Histograms show more detailed information for each metric (Fig. 4). For the number of voxels with ICC > 0.5 in short-term (session 2 against session 3) reliability, mPerAF had more voxels than mALFF and mfALFF (46336 vs. 44089 and 23148 voxels); zPerAF had more voxels than zALFF and zfALFF (46084 vs. 43510 and 22413 voxels) (Table 2). PerAF had fewer voxels than mPerAF and zPerAF (45122 vs. 46336 and 46084), but slightly more than mALFF and zALFF (45122 vs. 44089 and 43510), and much more than fALFF, mfALFF, and zfALFF (45122 vs. 26273, 23148, and 22413) (Table 2). For the long-term (session 1 against session 2) reliability, mPerAF had slightly more voxels than mALFF (31248 vs. 30866 voxels); and zPerAF also had slightly more voxels than zALFF (31743 vs. 31148 voxels). PerAF had fewer voxels than mPerAF, zPerAF, mALFF, and zALFF (22793 vs. 31248, 31743, 30866, 31148 voxels) (Table 2). The fALFF, mfALFF, and zfALFF showed the worst test-retest reliability (Table 2). Another long-term (session 1 against session 3) reliability showed similar results (Please see Fig. S2).

**Fig. 4.**
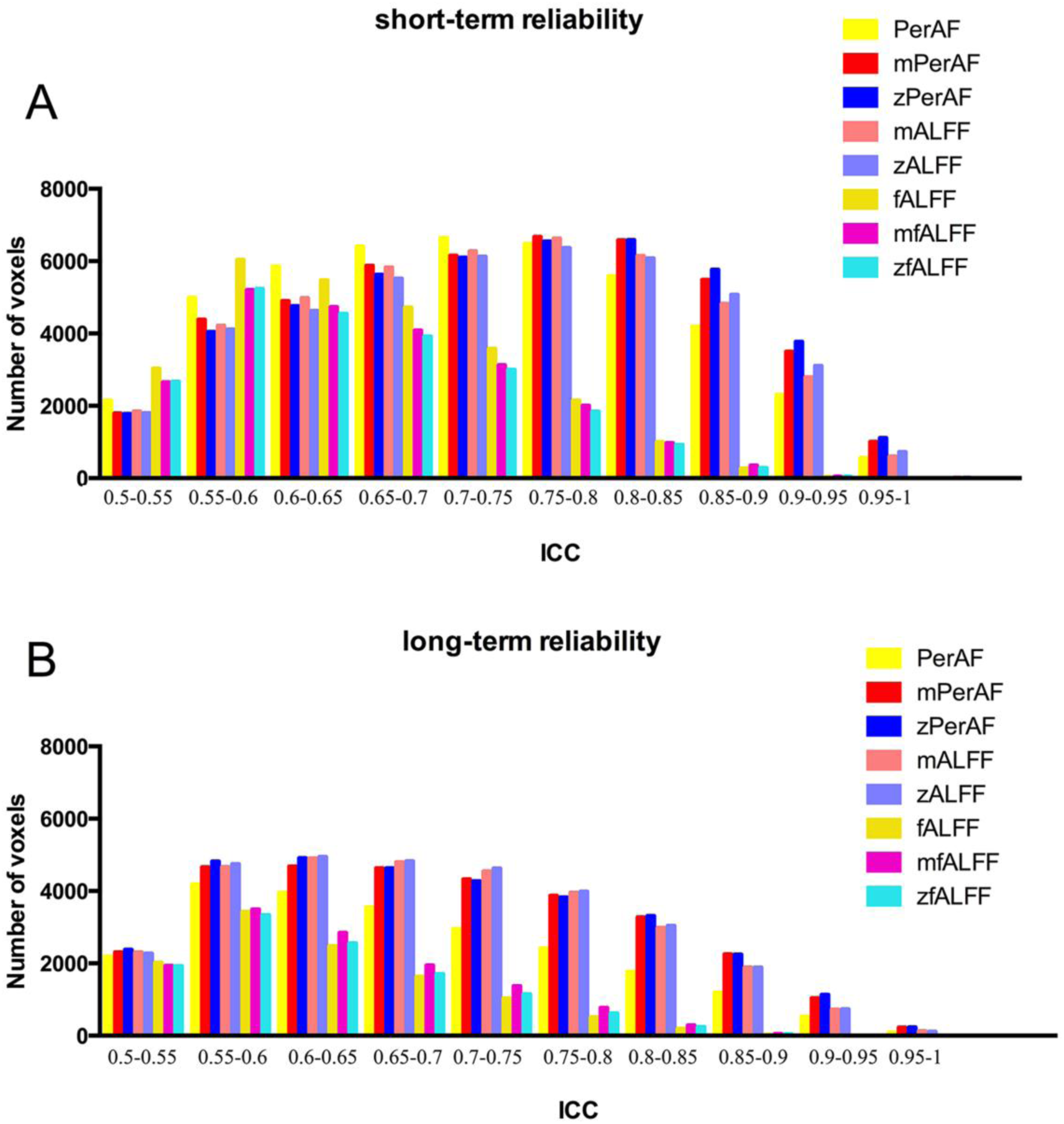
Histogram of test-retest reliability of all voxels. Y axis is the number of voxels of each bin (with an ICC step of 0.05). Upper (a) is the short-term (session 2 against session 3) reliability. In general, short-term reliability was better than long-term one. For short-term reliability, most voxels had ICC > 0.5 for all measures. Comparing the number of voxels with ICC > 0.5 among measures, PerAF, mPerAF, and zPerAF performed slightly better than mALFF and zALFF, and much better than fALFF, mfALFF, and zfALFF. Please see Table 2 for detailed number of voxels. For long-term reliability (session 1 against session 2), mPerAF and zPerAF performed similarly with (although very slightly better than) mALFF and zALFF, but PerAF (without standardization by global mean) performed worse, and fALFF, mfALFF, and zfALFF performed the worst.

**Table 2.** Number of voxels with ICC > 0.5

|  | Short-term reliability<br>(Session 2 against session 3) | Long-term reliability<br>(session 1 against session 2) |
| --- | --- | --- |
| PerAF | 45122 | 22793 |
| mPerAF | 46336 | 31248 |
| zPerAF | 46084 | 31743 |
| mALFF | 44089 | 30866 |
| zALFF | 43510 | 31148 |
| fALFF | 26273 | 11303 |
| mfALFF | 23148 | 12683 |
| zfALFF | 22413 | 11550 |

To view spatial overlap between PerAF and each of the other two methods, we selected the more comparable metrics, i.e., mPerAF, mALFF, and mfALFF, and performed overlapping analysis on voxels with ICC > 0.5 for all the metrics. As shown in Figure 5, mPerAF was largely overlapped with mALFF, and mfALFF was mostly included in mPerAF because the test-retest reliability of the mfALFF was lower than that of mPerAF and mALFF.

**Fig. 5.**
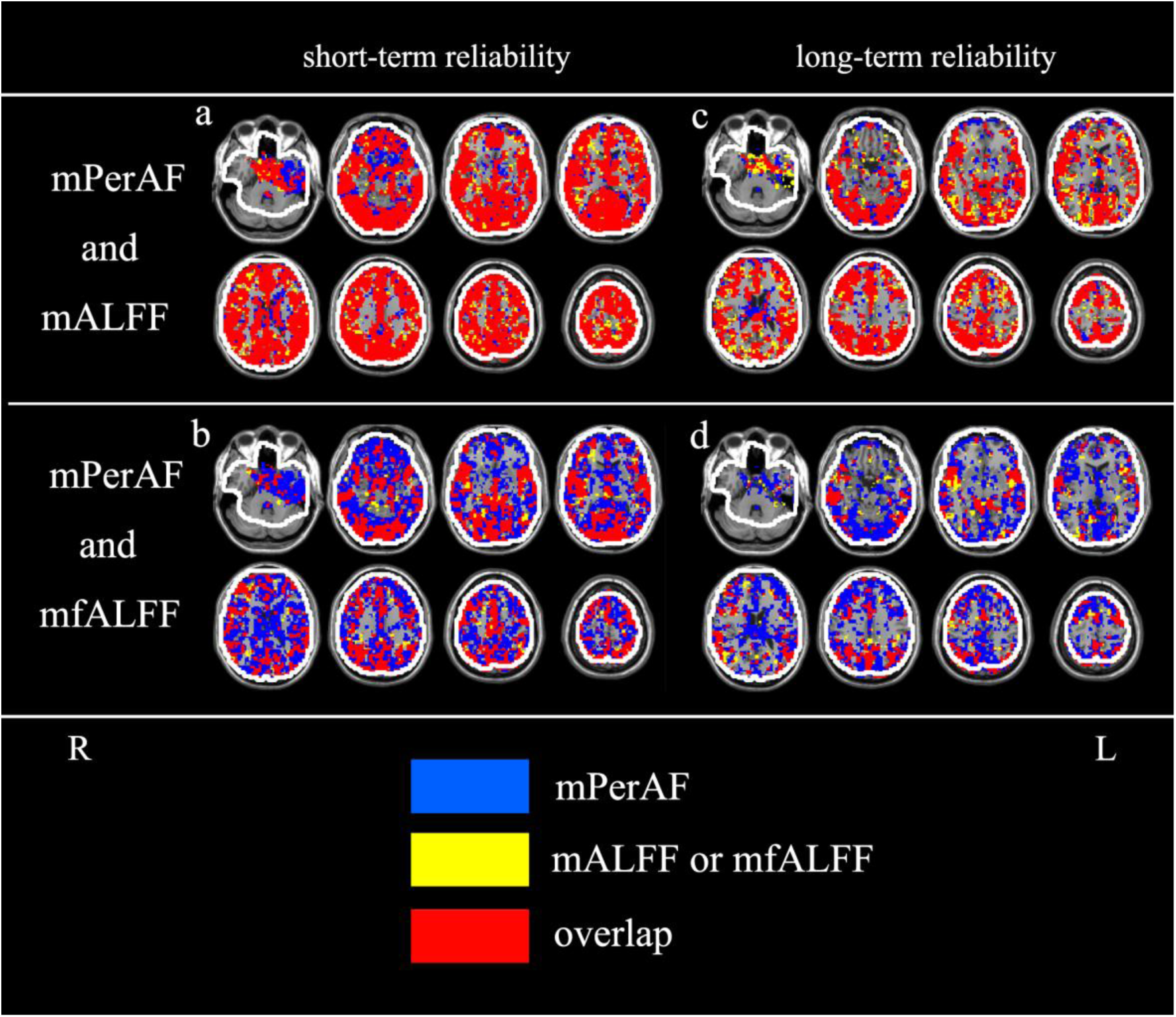
Test-retest reliability overlapping maps of mPerAF with mALFF and mfALFF on voxels with ICC > 0.5. The upper row is for mPerAF and mfALFF (a: short-term; b: long-term) and the lower row is for mPerAF and mfALFF (c: short-term; d: long-term). L: left side of the brain. R: right side of the brain.

Head motion regression slightly affect the ICC of both short- and long-term test-retest reliability (Fig. S3-5).

### 2.3. Dataset-2: comparison between EO and EC

#### 2.3.1. Data description

Dataset-2 was from a published data (Zou et al. 2015) which included 34 healthy participants (aged 19 - 31 years, 18 females). The study was approved by the Ethics Committee of the Center for Cognition and Brain Disorders, Hangzhou Normal University. Informed consent was obtained from each participant.

For each participant, four resting state sessions were scanned with two conditions EO and EC by BOLD and arterial spinlabeling (ASL), respectively. The order of the four sessions was counterbalanced across participants. The ASL data were not used in the current study. Dataset-2 was acquired using a GE MR-750 3.0 T scanner (GE Medical Systems, Waukesha, WI) at the Center for Cognition and Brain Disorders of Hangzhou Normal University. Each scan consisted of 240 contiguous EPI functional volumes (TR = 2000 ms; TE = 30 ms; flip angle = 60°; 37 axial slices; field of view (FOV) = 220 × 220 mm^2^, matrix = 64 × 64; in-plane resolution 3.44 × 3.44 × 3.4 mm^3^. For spatial normalization, a spoiled gradient-recalled pulse sequence was also used (176 sagittal slices; slice thickness = 1 mm; TR = 8100 ms; TE = 3.1 ms; flip angle = 9°; FOV=250 × 250 mm^2^).

#### 2.3.2. Data preprocessing

It was the same as mentioned in section “2.2.2”. In case not every participant’s whole brain was completely covered, we made an intersection mask within which all 68 scanning sessions (2 sessions for each of the 34 participants) were covered (Fig. 6). The detailed method was the same as in “2.2.2”. To compare the amount of head motion between EO and EC, we calculated framewise displacement head motion (Power et al. 2012). Framewise head motion calculates the relative head motion of each timepoint to its prior timepoint. Zang and colleagues used the sum of framewise head motion of ratation and transition separately (Zang et al. 2007) (See formuli 1 and 2 in that reference). Power and colleagues integrated the sum of 6 framewise headmtion parameters as a whole, named framewise displacement (FD) (Power et al. 2012). FD is beeing widely used and hence the current study also used FD (Power et al. 2012). Paired t-test was peformed on FD between EO and EC.

**Fig. 6.**
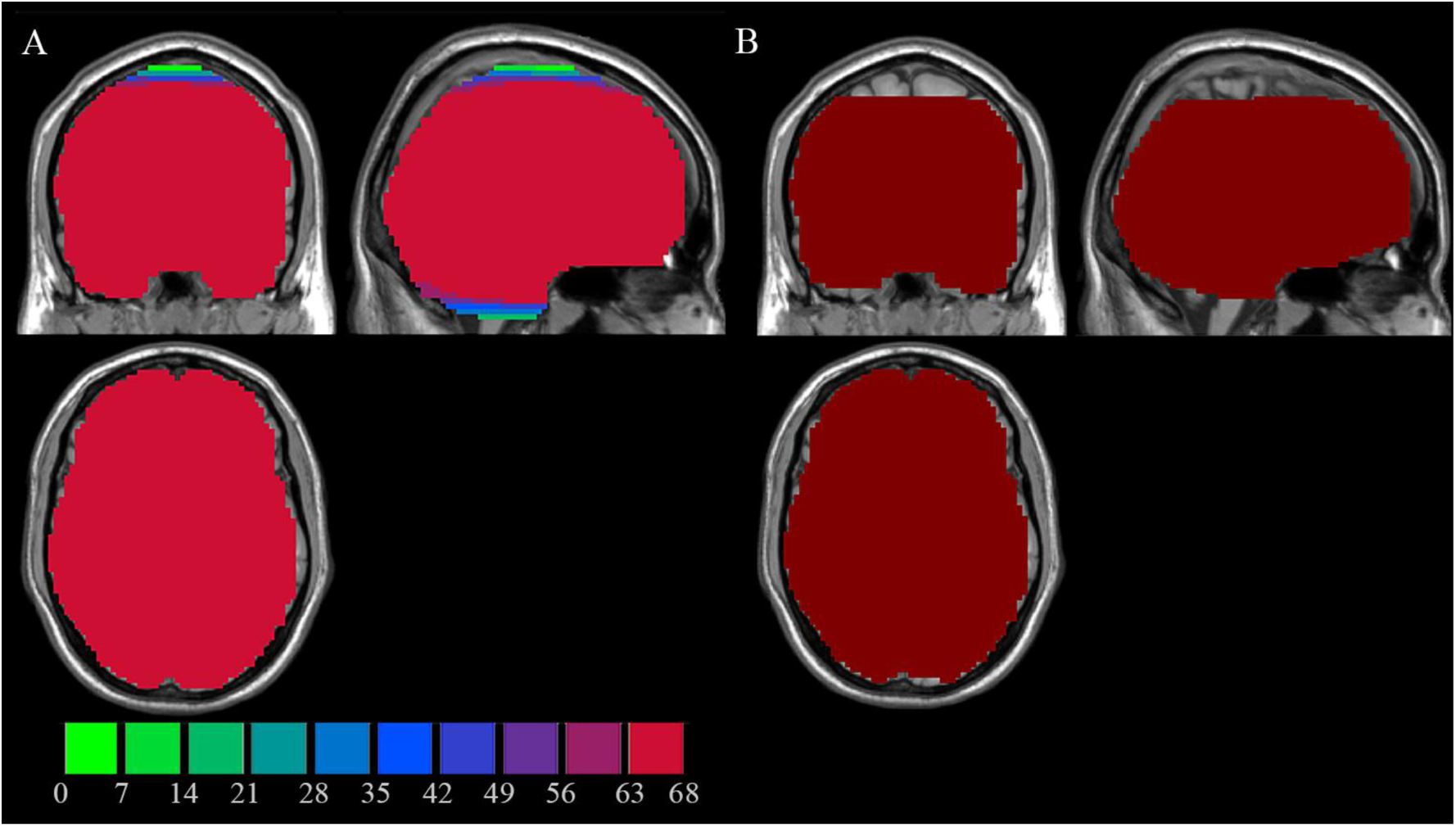
Intersection mask of Dataset-2. The left pannel shows how many sessions (totally 34 subjects × 2 = 68 session) were covered, for each voxel in Dataset-2. The right pannel is an intersection mask which was covered by all 68 sessions.

#### 2.3.3. Spatial pattern of PerAF

The calculation of PerAF, mPerAF, zPerAF, mALFF, zALFF, fALFF, mfALFF, and zfALFF was the same as those in Section 2.2.3.

To show the spatial pattern of PerAF, the averaged PerAF map of the 34 partipants in EC state was shown in Figure 7A. The pattern for EO was very similar with that of EC (not shown here). The averaged PerAF value of most voxels was from 0.14% (smallest) to 1.55% (< +2 SD) across the brain. Gray matter showed higher PerAF than white matter. The pattern for averaged fALFF, mPerAF, mfALFF, and mALFF were shown in Figure 7B, C, D, E, respectively. zPerAF, zfALFF, and zALFF were very similar with mPerAF, mfALFF, and mALFF, respectively, and not shwon here.

**Fig. 7.**
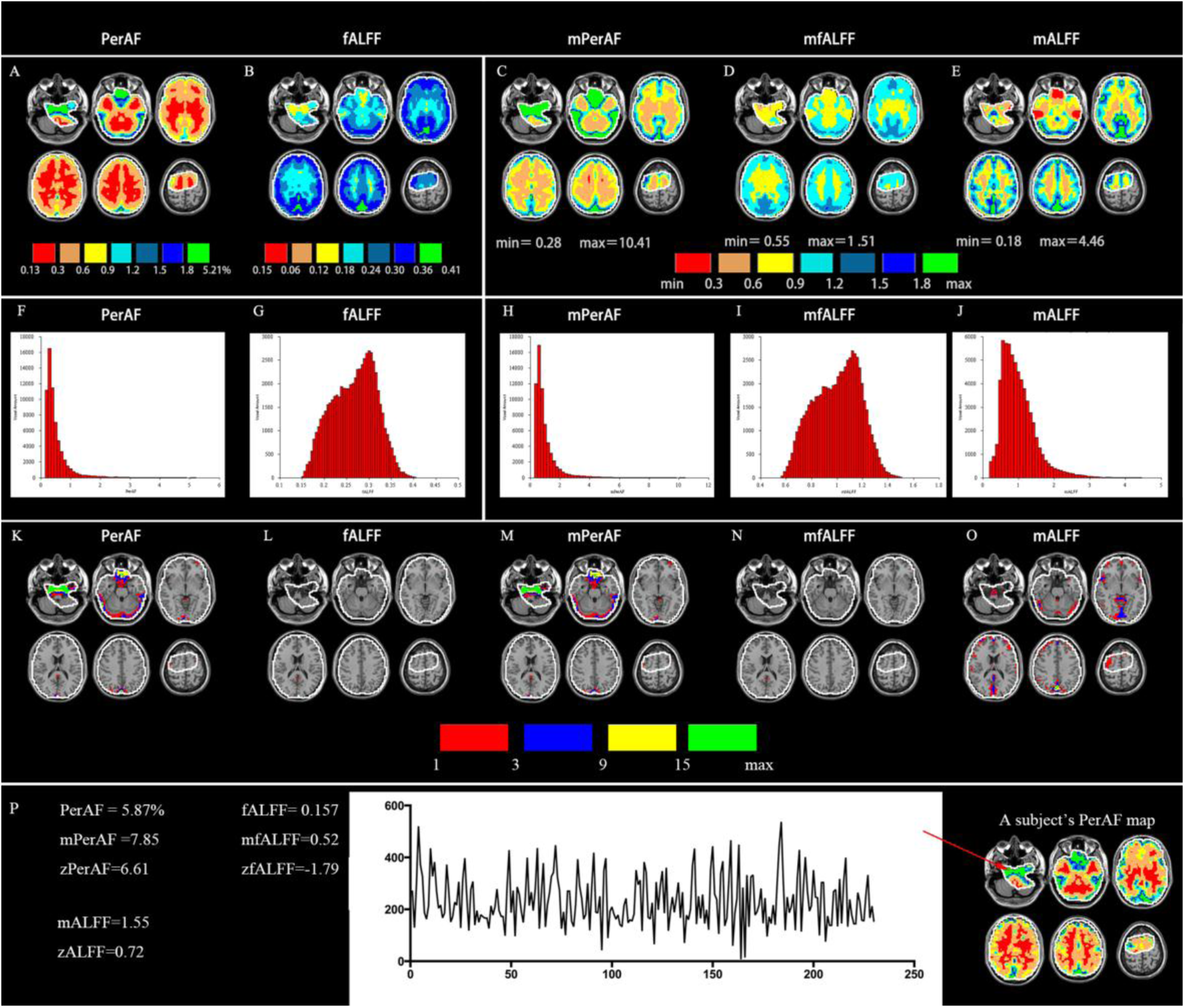
The spatial distribution of PerAF, ALFF, and fALFF as well as their derivatives. A – E: group averaged PerAF, fALFF, mPerAF, mfALFF, and mALFF (zPerAF, zfALFF, and zALFF were very similar with mPerAF, mfALFF, and mALFF, respectively and not shwon here). F – J: PerAF, fALFF, mPerAF, mfALFF, and mALFF, respectively (The histogram of zPerAF, zALFF, zfALFF was similar with mPerAF, mALFF, mfALFF, respectively, but no shown here. Please see Fig. S6). K – O: The pattern of extreme value (> 4 SD) of PerAF, fALFF, mPerAF, mfALFF, and mALFF. P: A timecourse of a participant’s voxel which showed very big PerAF (5.87 %). The BOLD signal intensity at some timepoints of this timecourse was nearly zero.

The histogram was quite different among the 3 measures (Fig. 7F-J). The histogram of averaged PerAF was very similar with that of averaged mPerAF; and averaged fALFF was similar with averaged mfALFF. The histogram of zPerAF, zALFF, zfALFF was similar with mPerAF, mALFF, mfALFF, respectively (Fig. S6).

The distribution as shown in the histogram of PerAF and mPerAF has a long tail at the right side (Fig. 7F, H). The distribution in the histogram of fALFF, mfALFF, and mALFF did show such long tail (Fig. 7G, I, J). The pattern of extreme value (> 4 SD) of PerAF, fALFF, mPerAF, mfALFF, and mALFF were different (Fig. 7K-O). For the PerAF and mPerAF, the voxels with extrmely high value were near the skull base (Fig. 7K). There was nearly no that big extreme value for fALFF and mfALFF (Fig. 7L, 7N). For mALFF, the voxels with extreme value were located either near large vessels and in the gray matter. We plotted a timecourse of a participant’s voxel which showed very big PerAF (5.87 %). As shown in Figure 7P, the BOLD signal intensity at some timepoints of this timecourse was nearly zero. No doubtly, this voxel has been affected by noise.

#### 2.3.4. Paired t-test between EO and EC for PerAF, ALFF, and fALFF

The calculation of PerAF, mPerAF, zPerAF, mALFF, zALFF, fALFF, mfALFF, and zfALFF was the same as those in section “2.2.3”.

Paired t-tests were performed between EO and EC. Multiple comparison correction was performed within the intersection mask. A combination of individual voxel’s P value < 0.05 and cluster size > 6156 mm^3^ was used, corresponding to a corrected P < 0.05 based on Monte Carlo simulation (rmm = 5, smothness = 6 mm, 1000 simulations) (from AFNI software and implemented in REST). In addition, to view potential differences between EO and EC outside the brain, the results of paired t-test for PerAF map (i.e., without standardization by global mean PerAF) was also shown without multiple comparison correction, i.e., only a voxel-level P < 0.05 without cluster size threshold was adopted.

In the case without standardization by global mean, significantly lower (corrected for multiple comparisons) PerAF in EO than in EC was observed in widely distributed brain regions including the bilateral primary sensorimotor cortex (PSMC), supplementary motor area (SMA), paracentral lobule, primary auditory cortex extending to superior and middle temporal gyrus, thalamus, precuneus, visual cortex, and posterior cingulate cortex (P < 0.05, corrected) (Fig. 8A1). Only small part of brain area (e.g., inferior orbital frontal, gyrus rectus) showed significantly higher PerAF in EO than EC.

**Fig. 8.**
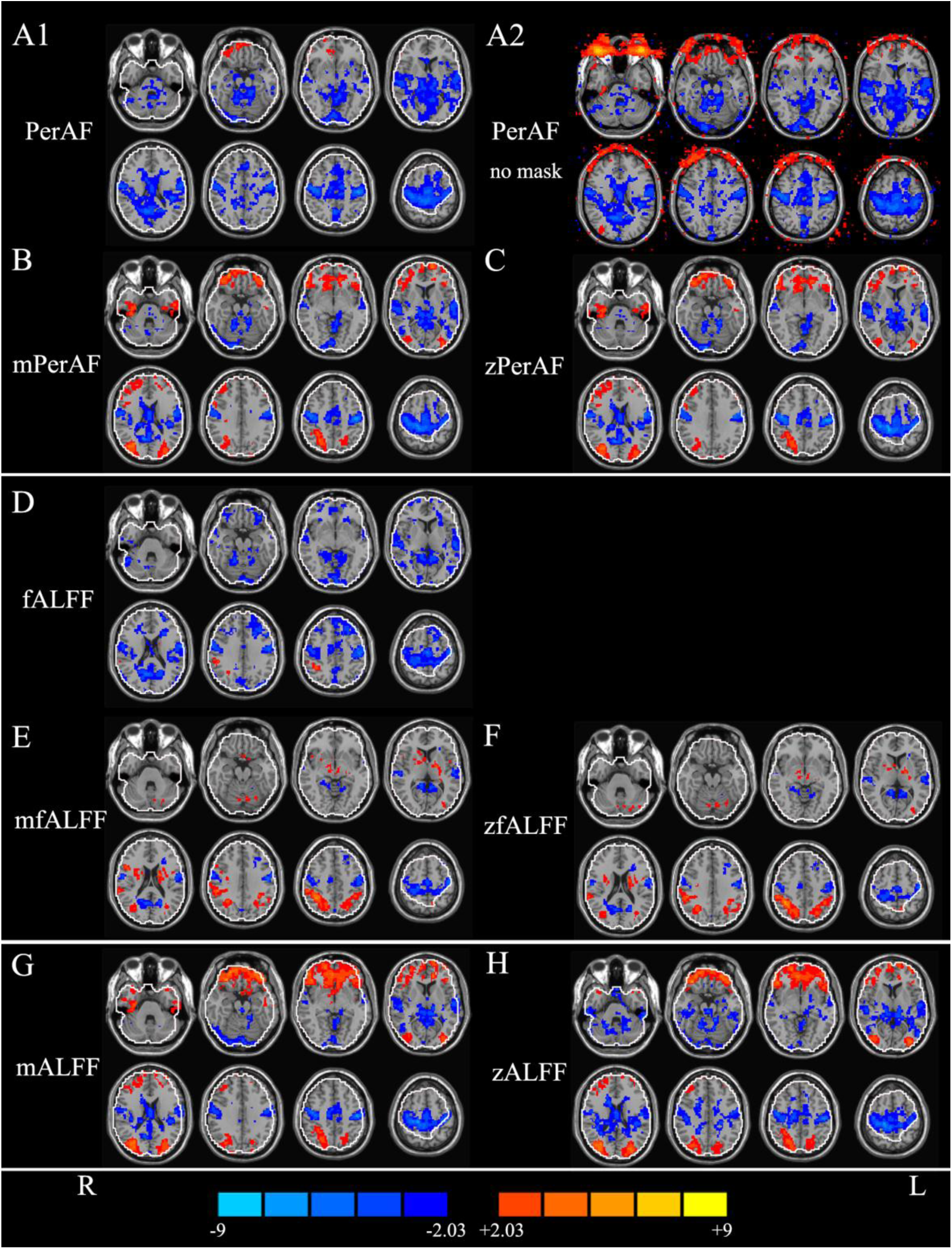
Results of paired t-test between eyes open (EO) and eyes closed (EC). A1: PerAF (without standardization by global mean) within brain intersection mask (P<0.05, corrected). A2: PerAF (without standardization by global mean) without brain mask (i.e., in the entire bounding box) (P<0.05, uncorrected). B: mPerAF (divided by global mean value) (P<0.05, corrected). C: zPerAF (minus mean then divided by standard deviation) (P<0.05, corrected). D - F: fALFF, mfALFF, and zfALFF, respectively (P<0.05, corrected). G, H: mALFF and zALFF, respectively (P<0.05, corrected). Warm colors indicate higher fluctuation in EO than EC, and cold colors indicate the opposite. L: left side of the brain. R: right side of the brain.

For fALFF (In the case without standardization by global mean), the pattern of difference between EO and EC was similar with that of PerAF, but with smaller volume for most clusters (Fig. Fig.8A1 vs. Fig.8D).

In the cases with global mean standardization, the between-condition difference of mPerAF (Fig. 8B), zPerAF (Fig. 8C), mALFF (Fig. 8G), zALFF (Fig. 8H) were very similar. Significantly higher fluctuation in EO than in EC was found in the bilateral middle occipital gyrus and orbitofrontal cortex. Significantly lower fluctuation in EO than in EC was found in the bilateral PSMC, SMA, paracentral lobule, thalamus, and primary auditory cortex (P < 0.05, corrected). For mfALFF (Fig. 8E) and zfALFF (Fig. 8F), the pattern of difference between EO and EC was generally similar with that of mPerAF, zPerAF, mALFF, and zALFF, except in the frontal pole and PSMC. mfALFF and zfALFF showed almost no difference in the frontal pole, while mPerAF, zPerAF, mALFF, and zALFF showed a big cluster. The cluster in the PSMC detected by mfALFF and zfALFF was smaller than that by mPerAF, zPerAF, mALFF, and zALFF.

The results of EO versus EC showed prominent inconsistency for comparisons with and without global mean standardization for PerAF (vs. mPerAF) as well for fALFF (vs. mfALFF) (Fig. 8A1 vs. Fig. 8B, Fig. 8D vs. Fig.8E). Specifically, in the case of no global mean standardization, only a small area showed higher fluctuation in EO than in EC. However, after standardization, a few other areas showed significantly higher fluctuation in EO than in EC, including the bilateral middle occipital gyrus and a large area in the orbitofrontal cortex. (not applicable for mfALFF and zfALFF results). Brain areas showing significantly lower fluctuation in EO than EC were slightly smaller than those without standardization. The prominent inconsistency suggested that the global mean PerAF had strong effect. We therefore performed a paired t-test on the global mean PerAF between the EO and EC. The global mean PerAF was calculated within a brain mask provided in REST (Song et al. 2011). It was found that the global mean PerAF was marginally lower in EO than EC (P = 0.0614).

When no brain mask was used and no multiple comparison correction was performed, the eyeballs showed significantly higher PerAF in EO than EC (Fig. 8A2). The difference in eyeballs extended to a large area of the frontal scalp and even to the orbitofrontal cortex.

The effect of head motion regression on the difference between EO and EC depended a lot on the measures used (Fig. 9). It showed very little effect on mPerAF, zPerAF, mALFF, and zALFF (Fig. 9 B, C, G, and H, respectively), but showed prominent effect on PerAF, fALFF, mfALFF, and zfALFF (Fig. 9A, D, E, F). Specifically, after Friston-24 head motion regression, the pattern of difference between EO and EC in PerAF was very similar with mPerAF (Fig. 9A-reg vs. Fig. 9B-no). Effects of Friston-24 head motion regression on fALFF, mfALFF, and zfALFF were quite interesting. Generally, a few clusters disappeared (Fig. 9 D, E, F).

**Fig. 9.**
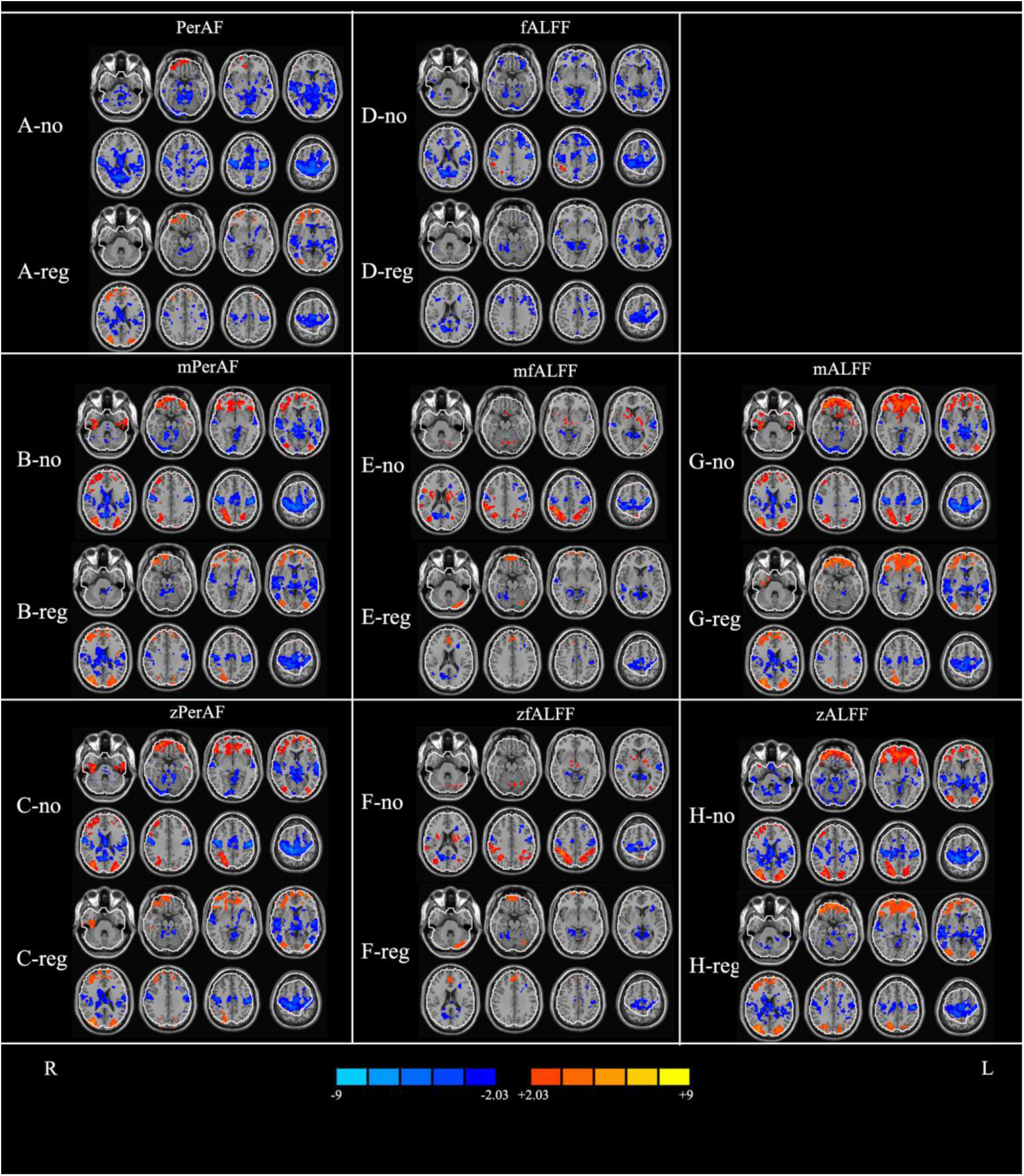
Effects of head motion regression (Friston-24 individually) on the results of paired t-test between eyes open (EO) and eyes closed (EC). –no: not regressed. –reg: regressed. All were corrected for multiple comparisons. Warm colors indicate higher fluctuation in EO than EC, and cold colors indicate the opposite. L: left side of the brain. R: right side of the brain.

One possible reason for the prominent effect of head motion on the PerAF difference between EO and EC might be due to potential difference of head motion between EO and EC. To test this assumption, we calculated the head motion amount, i.e., FD. The FD was 0.1036 ± 0.0331 (mean ± standard deviation) for EO, and 0.1095 ± 0.0514 for EC. There was no significant difference (P=0.3068).

## 3. Implementation and usage of PerAF calcuation toolkit

We developed a GUI toolkit, named REST-PerAF for PerAF calculation in MATLAB. It is an open source package based on some existing toolboxs including SPM (Friston et al. 1994) and REST (Song et al. 2011). From http://www.restfmri.net, the most recent version of REST-PerAF could be downloaded. The compressed package need to be extracted to a predefined directory, and then add the full path to MATLAB’s search path. PerAF is compatible with MATLAB version 7.1 or higher.

Entering ‘‘PerAF_GUI’’ in the MATLAB command window will open PerAF’s GUI. It supports NIfTI image format. Users need to set the input directory where the preprocessed data were in. The output directory also needs to be set. User can select PerAF (without standardization by global mean), mPerAF, or zPerAF.

We also implemented a command line in LINUX, named REST-PerAF, based on AFNI (Cox 1996) for calculation of PerAF. It can be downloaded at http://www.restfmri.net.

More usage details could be found in manual which can be downloaded at http://www.restfmri.net.

## 4. Discussion

As a measure of spontaneous brain activity, ALFF has been widely used in RS-fMRI studies (Margulies et al. 2010). However, as shown in the simulation in the current study (Fig. 1), ALFF is not a scale-independent measure. Therefore ALFF need to be standardized by the global mean value, producing mALFF or zALFF (Zang et al. 2007; Zuo et al. 2010a), or by the amplitude of entire frequency band (i.e., fALFF) (Zou et al. 2008). The calculation of mALFF and zALFF will face difficulties when the global mean value varies a lot (e.g., large lesions) or when only part brain is scanned due technical constraints (e.g., very fast imaging). fALFF will make the situation complex when taking sub-frequency bands into account. In this paper, we proposed PerAF as a single index for revealing the fluctuation of spontaneous activity at single voxel level.

### 4.1. Test-retest reliability

As shown in Fig. 3, all measures including PerAF, mPerAF, zPerAF, mALFF, zALFF, fALFF, mfALFF, and zfALFF showed moderate to higher short-term test-retest reliability, but the long-term reliability was not so good. These results were consistent with previous studies on the same dataset by using zALFF and zfALFF (Zuo et al. 2010a). For both short- and long-term reliability (Fig. 4A and Fig. 4B, Table 2), mPerAF and zPerAF showed slightly higher ICC than mALFF and zALFF, respectively. This slightly better performance may be partly contributed by normalizing the temporal mean of BOLD signal of PerAF. However PerAF had less number of voxels with ICC > 0.5 than mPerAF and than zPerAF, suggesting that further standardization by global mean could improve test-retest reliability. Anyway, it does not mean that mPerAF or zPerAF could replace PerAF, as discussed in the following session of between-condition differences.

The test-retest reliability of fALFF, mfALFF, and zfALFF were much worse than PerAF, mPerAF, zPerAF, mALFF, and zALFF. These results were well consistent with the study by Zuo and colleagues (Zuo et al. 2010a). The bad performance of fALFF is probably due that it takes high frequency band into account.

### 4.2. Between-condition differences with and without standardization by global mean value

One of the advantages of PerAF is that it is not necessarily to be standardized by global mean value and can be directly used for statistical analysis. However, it is interesting that with and without standardization yielded quite different results. Without standardization, the areas showing significant differences in the bilateral PSMC, SMA, primary auditory cortex, and primary visual cortex became smaller (Fig. 8A1). But after standardization, these clusters became smaller and two more regions, i.e., the lateral occipital cortex and orbitofrontal cortex, appeared (Fig. 8B, C). The lateral occipital cortex plays important role in visual information processing, and the difference of fluctuation between resting EO and EC is reasonable. As PerAF, fALFF can also be used without standardization by global mean value. The global mean fALFF also showed prominent effect on the results (Fig. 8D vs. E and F). These results suggest that, standardization by the global mean value has prominent effect on the results, and both results seem meaningful. We further compared the global mean PerAF between EO and EC resting states, a marginally higher PerAF was found in EC than EO resting states (P=0.0614). Jao and colleagues also found that there were some global differences of fluctuation amplitude between EO and EC (Jao et al. 2013).

Taken together, although the test-retest reliability of mPerAF and zPerAF is better than PerAF, results of both with and without removing global mean PerAF seemed to be physiologically meaningful. The impact of global mean normalization could not be neglected. However, its actually physiological or psychological importance is still unclear and needs further investigation.

In addition, the results of two ways of standardization, i.e., z-transformation and dividing global mean, were almost the same (Fig. 8 B vs. C, E vs. F, G vs. H). And actually, the histogram of dividing global mean value and z-transformation were also very similar (Fig.7 H-J vs. Fig.S6 A-C).

### 4.3. Artifacts from eyeball

Without global mean standardization, the difference between EO and EC in the orbitofrontal cortex was much less than that after global mean standardization (Fig. 8A1 and Fig. 8D vs. Fig.8 B, C, E, F, G, H). As shown in Fig. 8A2, the eyeballs showed significant higher PerAF in EO condition, and this difference extended to frontal scalp and even to orbitofrontal cortex. Therefore, the prominent difference in the orbitofrontal cortex was probably an artifact of eyelid movements during EO condition, which might have resulted in a change of magnetization in the adjacent area. Further, the global mean standardization magnified this artifact in the orbitofrontal cortex. A recent study from our group used fast sampling (TR = 400 ms) and compared the amplitude of RS-fMRI between EO and EC in different sub-frequency bands (Yuan et al., 2014). The difference in the orbitofrontal cortex was found only in higher frequency band (> 0.1 Hz), but almost no significant difference was found within 0.01 – 0.08 Hz. Further, such difference seemed to be more prominent from 0.4 Hz to 1.25 Hz (with a step of 0.05 Hz). Such a high frequency difference is unlikely to be consequence of neuronal activity within the brain. Unfortunately, our previous study did not cover the eyeball area due to technical limitation of fast sampling (only 13 slices were scanned), therefore, we failed to investigate the higher frequency amplitude difference between EO and EC in the eyeball area (Yuan et al., 2014). Future studies can record the eye blinking by electro-oculogram or video camera and use fast sampling, then quantitatively analyze the effect of eye blinking on the PerAF in the orbitofrontal cortex.

### 4.4. Effects of head motion regression

In-scanner head motion (Van Dijk et al. 2012) has been widely taken as nuisance or artifact in RS-fMRI studies. While most studies have proposed that head motion has prominent effect on functional connectivity (e.g., Van Dijk et al., 2012), a few studies reported that head motion also had prominent effect on ALFF or fALFF, two similar methods for measuring single voxel activity (Satterthwaite et al. 2012; Yan et al. 2013a). The current study aimed to investigate the head motion effect on PerAF, ALFF, fALFF, as well as their derivatives. It should be noted that, rather than the effect on a single resting condition, we evaluated the head motion effect on the difference between two distinct RS-fMRI conditions, eyes open (EO) and eyes closed (EC). The difference of ALFF between EO and EC in the human brain has been well documented with independent datasets (Yang et al. 2007; Liu et al. 2013; Jao et al. 2013; Yuan et al. 2014; Liang et al. 2014; Zou et al. 2015). As recommended in the study by Yan and colleagues, we also applied Friston-24 head motion parameters to regress out head motion individually. We found that, the head motion effect on the difference between EO and EC was prominent for PerAF, fALFF, mfALFF, and zfALFF, but very little for mPerAF, zPerAF, mALFF, and zALFF (Fig. 9). Interestingly, the head motion effect for PerAF was similar with the effect of global normalization (i.e., PerAF vs. mPerAF and zPerAF) (Please see Fig. 9A-reg vs Fig. 9B-no, and Fig. 9A-reg, vs Fig. 9C-no). This phenomenon was, to some extent, consistent with a notion that global timecourse removal effectively reduces motion-related functional connectivity (Yan et al. 2013a). Like the debates on global timecourse removal for functional connectivity analysis, head motion is not always artifact or nuisance. For example, a few studies have reported that head motion was related to impulsivity in a large sample of healthy participants (Kong et al. 2014), to heritability in a twin study (Couvy-Duchesne et al. 2014), and to stereotype of head rotation symptom of Down Syndrome (Pujol et al. 2014). Therefore, we suggest that, while individual head motion regression is a necessary procedure during RS-fMRI preprocessing, results of both with and without head motion regression should be presented.

The head motion effect on fALFF, mfALFF, and zfALFF seems a little complex and more prominent than that on PerAF, mPerAF, zPerAF, mALFF, and zALFF. Generally, a few clusters disappeared (Fig. 9D-reg, Fig.9E-reg, Fig.9F-reg). The result of more prominent effect on fALFF than PerAF and ALFF was consistent with the results found by Satterthwaite and colleagues (Satterthwaite et al. 2012). We agree with their interpretation that head motion may introduce high-frequency artifact into the data and hence more effect on fALFF (Satterthwaite et al. 2012) because fALFF algorithm takes high frequency signal into account (Zou et al. 2008).

### 4.5. Limitations

As shown in Figure 7P, PerAF was sensitive to a temporal noise. Temporal signal to noise ratio (tSNR) is a widely used measure in fMRI studies and is an inverse of PerAF. We did not scan phantom data to measure tSNR. Nor did we measure spatial SNR. Both temporal and spatial SNR can be affected by multiple factors, e.g., voxel resolution, TR, TE, FA, acceleration factor (iPAT) and many others (Triantafyllou et al. 2011).

We proposed that it is not necessary for PerAF to be standardized by global mean, and hence is more suitable for brain lesion studies than ALFF. However, we didn’t include any data with lesions. It should be further evaluated how big effect the brain tumor or other large lesions have on global mean normalization.

## 5. Conclusions

PerAF is an analog to percent signal change widely used in task fMRI studies, and hence a straightforward measure of BOLD signal fluctuations at single voxel level. The test-retest reliability showed that PerAF was generally slightly higher than conventional ALFF, and much better than fALFF. With and without standardization of global mean PerAF yielded quite different results, suggesting that with and without global mean standardization are not exclusive. Both results of test-retest reliability and between-condition differences suggested that PerAF is a more promising measure for RS-fMRI signal at single voxel level.

## Information Sharing Statement

Dataset-1 in this article is freely available from http://www.nitrc.org/projects/fcon_1000/.

## Acknowledgements

This study was supported by grants from the National Natural Science Foundation of China (81520108016, 81271652, 81020108022, 31471084, 81471653, 81201155, 81301210, 81201156, 81401400, 81201083). Ze Wang is supported by the State Youth 1000 Talent Program of China and Hangzhou Qian Jiang Endowed Professorship. Dr. Zang is partly supported by “Qian Jiang Distinguished Professor” program. The authors thank Mr. YUAN Bin-Ke for his help on data collection.

